# HiIDDD: A high-throughput imaging pipeline for the quantitative detection of DNA damage in primary human immune cells

**DOI:** 10.1101/2021.10.20.465132

**Authors:** Kelsey Gallant, Arsun Bektas, Mary Kaileh, Ana Lustig, Luigi Ferrucci, Gianluca Pegoraro, Tom Misteli

## Abstract

DNA damage is a prominent biomarker for numerous diseases, including cancer and aging. Detection of DNA damage routinely relies on traditional microscopy or cytometric methods. However, these techniques are typically of limited throughput and are not ideally suited for large-scale longitudinal and population studies that require analysis of large sample sets. We have developed HiIDDD (<u>Hi</u>gh-throughput Immune cell <u>D</u>NA <u>D</u>amage <u>D</u>etection), a robust, semiquantitative and single-cell assay that measures DNA damage by high-throughput imaging using the two major DNA damage markers 53BP1 and γ−H2AX. We demonstrate sensitive detection of DNA damage in a wide set of freshly isolated and cryopreserved primary human immune cells, including CD4^+^ and CD8^+^ T cells, B cells and monocytes with low inter-assay variability. As proof of principle, we demonstrate parallel batch processing of several immune cell types from multiple donors. We find common patterns of DNA damage in multiple immune cell types of donors of varying ages, suggesting that immune cell properties are specific to individuals. These results establish a novel high-throughput assay for the evaluation of DNA damage in large-scale studies.

## Introduction

A large number of DNA lesions occur daily in a human cell. DNA damage is caused by exogenous triggers, such as cytotoxic chemicals or irradiation, and by endogenous events such as DNA replication^1–3^. Cells counteract DNA damage through several DNA damage response (DDR) pathways that are activated based on the type of lesion present^1,2,4,5^. Homologous recombination (HR) uses sister chromatids as a template for repair and acts both on single- and double strand breaks (DSBs) and is limited to the S- and G2-phase of the cell cycle. The majority of DSBs, however, are eliminated by non-homologous end joining (NHEJ), in which DSBs are repaired by ligation of the gap without use of a template, often leading to short deletions or insertions^6^.

NHEJ requires a complex protein machinery to execute the repair process^7,8^. In human cells the core histone variant H2AX is phosphorylated at Ser139 (γ-H2AX) in response to DNA damage^9^. Once established, γ-H2AX acts as a binding site for various repair factors that ultimately execute the repair reaction^7,8^. One prominent NHEJ repair factor is the tumor suppressor p53-binding protein 1 (53BP1), which colocalizes with □-H2AX and promotes NHEJ-mediated repair^10,11^.The accumulation of □-H2AX and 53BP1 is detected as distinct foci in the nucleus of cells that contain DNA damage and can be visualized with high reliability by indirect immunofluorescence using specific antibodies^7,10–12^. Because of their prominent involvement in NHEJ and robust detection, both 53BP1 and □-H2AX have been widely used as biomarkers of DSBs and of DDR activity^12–15^.

DNA damage is a physiologically relevant biomarker for various diseases and pathological conditions, including cancer and inflammation, and it is routinely used in the assessment of immunotherapy effects, drug action, and radiation exposure^16–18^. Determination of DNA damage is of particular interest in monitoring human aging since genome instability and increased DNA damage have been proposed to contribute to the aging process^19–21^. In support of this view, many premature aging disorders, such as Werner Syndrome, Cockayne Syndrome or Hutchinson Gilford Progeria Syndrome, are characterized by increased DNA damage^22,23^. Moreover, an elevated mutational load in DNA repair-associated genes has been linked to numerous aging-related cancers^17,24,25^. However, the extent to which DNA repair capacity contributes to the normal aging process remains unclear.

Standard methods to quantitatively measure DNA damage accumulation and DNA repair capacity in human cells are well established and include use of alkaline comet assays, indirect fluorescence microscopy using antibodies and flow cytometry^5,26–31^. However, these assays are often limited in that they either measure bulk DNA damage, as in the case of comet assays or are of low throughput, such as microscopy and cytometry methods. These limitations make these assays unsuitable to probe large numbers of primary human samples that would be needed for comprehensive longitudinal or epidemiological studies, and to assess the role of DNA damage in many physiological settings.

We have developed here a versatile assay for high-throughput detection of DNA damage in multiple primary immune cell types. The assay is based on the detection of the two prominent DNA damage markers □-H2AX and 53BP1 using a high-throughput imaging platform. Peripheral blood mononuclear cells (PBMCs) are a suitable tissue source for large-scale studies since they can be easily collected, and since they serve as a sensitive and global indicator of an individual’s physiological status. However, immune cells and PBMCs tend to be a challenge in microscopy-based approaches due to their fragility and poor adherence to substrates used in most imaging methods^32,33^. We have overcome these limitations by optimizing an assay, HiIDDD (**Hi**gh-throughput **I**mmune cell **D**NA **D**amage **D**etection), which uses immunofluorescence (IF) and high-throughput imaging (HTI), combining automated liquid handling, microscopy, and image analysis (Fig. 1a). This approach allows imaging and quantitative analysis of up to tens of thousands of individual cells in a large numbers of samples in a single experiment^34,35^. In HiIDDD, we have optimized cell seeding, fixation, permeabilization, and immunostaining steps to create a robust HTI IF protocol for detection of DNA damage on immobilized T cells, B cells, and monocytes in 384-well microplates. We demonstrate that this approach is a sensitive and reliable assay for the measurement of DNA damage in primary human immune cells.

**Fig. 1).**
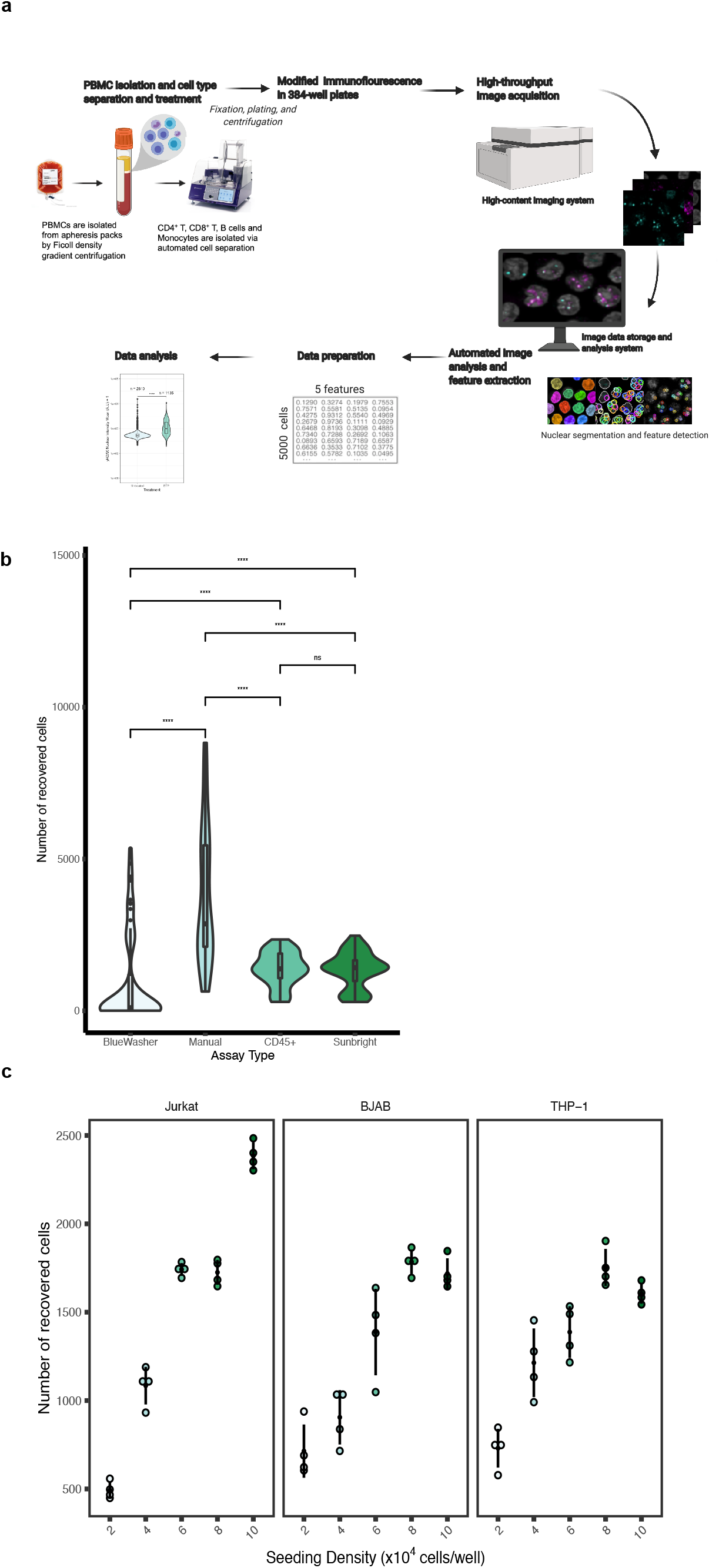
High-throughput imaging of DNA damage in human immune cells. **a)** Schematic representation of the high-throughput imaging methodology to detect DNA damage in CD4^+^T, CD8^+^T, B cells and monocytes. The pipeline involves a modified immunofluorescence assay, imaging using an automated high-content screening system, image analysis for segmentation and fluorescence intensity cellular features extraction, and statistical analysis. **b**) Determination of seeding efficiency using DAPI staining ∼15 hours after seeding. Additional substrates were added to poly-D-lysine plates; cells were seeded using the BlueWasher liquid handler unless indicated otherwise. Values represent at least 48 replicate wells per condition; box represents first and third quartiles. **c)** Jurkat, BJAB, and THP-1 cells were seeded at densities ranging from 2-10×10^4^ in a 384-well format. The total number of detected nuclei by DAPI is indicated. N= 4 wells per condition with at least 467 cells per condition.

## Results

### Detection of DNA damage in immune cell lines in 384-well format

We sought to establish a high-throughput imaging pipeline as a rapid and robust method for the detection and measurement of DNA damage and repair capacity in immune cells. The HiIDDD pipeline consists of plating of immune cells in 384-well plates, detection of □-H2AX and 53BP1 by IF, automated imaging, image analysis, and statistical analysis (Fig. 1a). For HiIDDD development, a set of semi-automated steps for sample preparation, indirect immunofluorescence (IF) staining, and washing methods were optimized to maximize the adherence of a range of immune cell types and to minimize technical variability between replicates (Fig. 1; see Materials and Methods). Primary antibodies against two prominent biomarkers of DNA damage and repair, 53BP1 and □-H2AX, were used based on their previously demonstrated efficiency in IF staining protocols for multiple cell types. The assay was designed to be amenable to a wide range of high-content data acquisition and analyses parameters for cellular features detected (e.g., foci number, foci intensity, nuclear intensity, etc.).

Most immune cells grow in suspension and are notoriously difficult to immobilize on a glass or plastic surface, which is generally necessary for imaging. To optimize the immobilization of immune cells on a substrate for HTI with automated liquid handling, we explored use of various surface coatings including large surface glycoproteins (e.g. CD45 and CD43) or activated polyethylene glycol (e.g. SUNBRIGHT®) previously shown to promote immune cell attachment^36^, but found only a modest increase in adhesion compared to routinely used poly-D-lysine coating alone (Fig. 1b). We thus first optimized cell seeding density and evaluated adherence of Jurkat, BJAB and THP-1 cell lines, representing T cells, B cells and monocytes, respectively, on poly-D-lysine coated 384-well plates (see Materials and Methods). To determine optimal cell seeding density, cells were plated in 384-well plates in a range from 2×10^4^ cells/well to 1×10^5^ cells/well in a total volume of 40 µl per well (Fig. 1c). Cells were fixed and plates were centrifuged briefly (400x g, 4 min, RT) and post-fixed in 4% paraformaldehyde for 20 min at RT. Importantly, single fixation before or after centrifugation of cells and use of a robotic dispenser resulted in significantly lower cell yields (Fig. 1b). Using manual dispensing onto poly-D-lysine coated wells, seeding density was evaluated based on the number of recovered cells detected by DAPI and variability between four technical replicates for each density after staining for □-H2AX or 53BP1. The optimal seeding density was determined as ∼0.6-1.0×10^5^ cells/well for Jurkat and ∼8×10^4^ cells/well for BJAB and THP-1 cells (Fig. 1c).

### Detection of DNA damage by HTI in immune cell lines

In order to determine whether an increase in DNA damage can be detected by HTI, Jurkat, THP-1, and BJAB cells were treated with 30 µM of etoposide (ETP) for 1.5 hrs to induce dsDNA breaks (DSB) and cells were subjected to HTI immunostaining for 53BP1 and □-H2AX (Fig. 2a). For each condition, three technical replicate wells were imaged in nine randomly selected fields of view (FOV) per well, corresponding to approximately 1-5×10^3^ cells per condition, and single-cell signal intensity for 53BP1 and □-H2AX was measured using automated image analysis (see Materials and Methods). Typically, 2,000-5,000 cells were analyzed per sample. Distinct fluorescence intensity parameters were used for the two markers based on their different changes in nuclear fluorescence distribution in response to DNA damage in the primary immune cells used here. For 53BP1, which is present in the cell nucleus prior to DNA damage but accumulates in distinct foci upon DNA damage, integrated spot intensity was measured, whereas for □-H2AX, which is present at low levels in undamaged cells and rapidly increases upon DNA damage due to phosphorylation at S139^ref.9^ total mean nuclear intensity was measured^10,11^. As expected, upon ETP treatment 53BP1 and □-H2AX rapidly formed nuclear foci at sites of DNA damage (Fig. 2a) and we observed an increase in both 53BP1 and □-H2AX pan nuclear staining, defined as the detected fluorescence of □-H2AX within segmented nuclei (Fig. 2a).

**Fig. 2).**
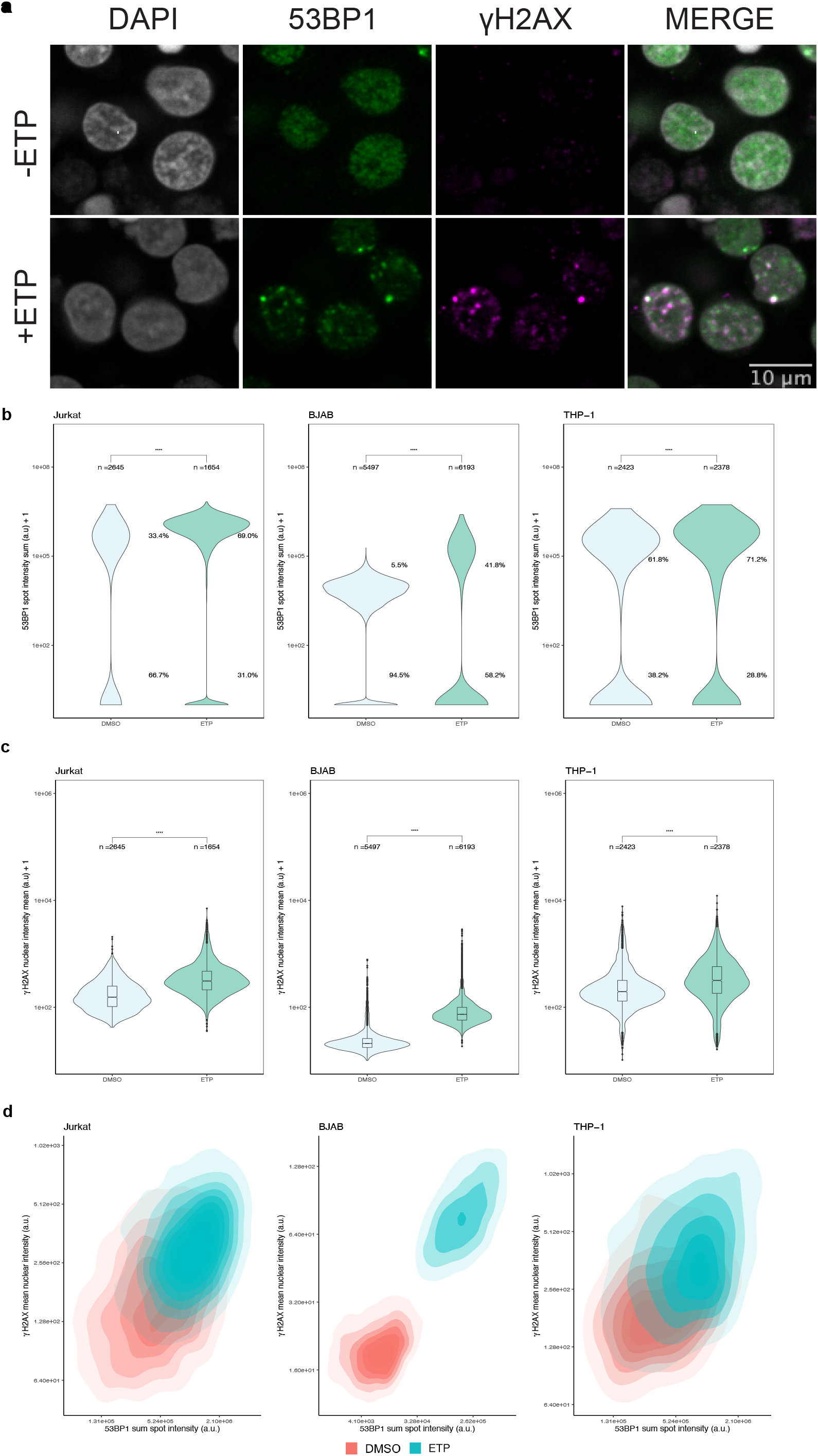
Detection of DNA damage in immune cell lines using HTI. **a)** Jurkat cells were either treated with DMSO or treated with 30 µM etoposide (ETP) for 1.5 hrs before being subjected to HTI. DNA damage sites are indicated by an increase in signal intensity and foci formation of 53BP1 and □-H2AX detected by specific antibodies. Scale bar: 8µm. **b)** Distribution of 53BP1 sum spot intensity in DMSO- or ETP-treated cells. ****: p < 0.0001. **c)** Distribution of *γ*-H2AX mean nuclear intensity in DMSO or ETP-treated immune cells. ****: p <0.0001 using Mann-Whitney-Wilcoxon test. Number of analyzed cells (n) indicated for each sample. The values are representative of at least 3 independent experiments. Boxes represent the first and third quartiles of the distribution, the line represents the median and the whiskers represent the highest and lowest data points within 1.5 times the interquartile range (IQR) above the upper quartile and below the lower quartile. **d)** Two-dimensional kernel density contour plots of the correlation between 53BP1 sum spot intensity and □-H2AX mean nuclear intensity for both ETP-treated (blue) and untreated (red) immune cell lines for the dataset shown in panels B and C. N = 3 wells per sample with at least 197 cells per well.

In Jurkat cells, integrated 53BP1 spot intensity per cell increased significantly upon DNA damage by ETP (median: 1.06 ± 0.93 e6 A.U.; standard deviation) compared to vehicle control (median: 4.67 SD: ± 5.85 e5 A.U.) (Wilcoxon test p-value *<* 2.2 e-16) (Fig. 2b). Importantly, using single cell analysis we observed an increase in the number of cells that were 53BP1 positive when treated with ETP compared to the vehicle treatment (69.0% versus 33.4%; *X*^2^ = 518.49, *p <* 2.2 e-16; Supplementary Fig. 1) when applying a threshold of one standard deviation (SD) from the mean of the negative control. Likewise, ETP-treated Jurkat cells (median: 432 ± 459 A.U) showed a significant increase in □-H2AX mean nuclear intensity when compared to the vehicle control (median: 199 ± 151 A.U) (Fig. 2c; Wilcoxon test p-value p *<* 2.2 e-16). At the single cell level, □-H2AX was detected in 42.3% of ETP-treated cells compared to 12.4% in untreated controls cells using as a threshold one standard deviation from the mean □-H2AX nuclear intensity of the negative control (Fig. 2c; Supplementary Fig. 1; *X*^2^ = 499.09, *p <* 2.2 e-16; DMSO: n= 2645 ETP: n = 1654).

Similar results were found for BJAB and THP-1 cells (Fig. 2b). For BJAB cells, integrated 53BP1 spot intensity per cell increased about 5-fold from 3.21 ± 0.71 e4 A.U. in controls to 1.78 ± 1.57 e5 A.U. in ETP-treated cells (Fig. 2b, Wilcoxon test p-value *<* 2.2 e-16). In addition, 41.8% of cells were 53BP1-positive when compared to the vehicle treatment (5.4%) (*X*^2^ = 2057.2, p < 2.2 e-16). □-H2AX mean nuclear intensity increased from 20.1 ± 5.79 A.U. in controls to 72.8 ± 28.1 A.U. in ETP-treated samples (Fig. 2c, Supplementary Fig. 1; middle, Wilcoxon test p-value < 2.2 e-16). At the single cell level, □-H2AX was detected in 75.9% of ETP-treated cells compared to 5.7% in untreated controls cells (Fig. 2c, Supplementary Fig. 1; *X*^2^ = 5855.1, p < 2.2 e-16; DMSO: n = 5497 ETP: n = 6193).

For THP-1 monocytes, we found a significant increase in integrated 53BP1 spot intensity per cell in response to ETP-induced DNA lesion (median: 6.25 ± 6.07 e5 A.U.) as compared to vehicle control (median: 3.37 ± 2.95 e5 A.U.) (Wilcoxon test p-value < 2.2 e-16) (Fig. 2b). Similarly, 32.4% of cells were 53BP1 positive when compared to the vehicle treatment (13.6%) (*X*^2^ = 177.36, p < 2.2 e-16). As expected, there was also an increase in □-H2AX mean nuclear intensity (Fig. 2c, Supplementary Fig. 1; Wilcoxon test p-value < 2.2 e-16) when comparing ETP-treated cells (median: 317 ± 250 A.U.) to vehicle control (median: 196 ± 117 A.U.). At the single cell level, □-H2AX was detected in 9.46% of ETP-treated cells compared to 5.65% in untreated controls cells using as a threshold of one SD from the mean of □-H2AX mean nuclear intensity in the negative control (Fig. 2c, Supplementary Fig. 1; *X*^2^ = 3.71, p = 0.054; DMSO: n = 1591 DMSO: n= 2348).

To relate the behavior of the two DNA damage markers to each other and to ask whether both markers simultaneously increased in individual cells in response to ETP, or if they represent cell-autonomous events, integrated 53BP1 spot intensity per cell and □-H2AX mean nuclear intensity per cell were plotted as two-dimensional kernel density estimates (Fig. 2d). ETP treatment shifted the density distributions to higher values of both parameters in all cell types compared to untreated conditions, suggesting concomitant responses in most cells in the population (Fig. 2d).

### Detection and measurement of 53BP1 and □-H2AX in primary immune cells using HTI

To assess whether HiIDDD can detect and quantitatively measure DNA damage in primary immune cells, we applied the pipeline to freshly isolated and frozen CD4^+^T cells, CD8^+^T cells, B cells and monocytes. Similar to cultured cell lines, and following ETP treatment, isolated immune cells were immediately fixed in 4% PFA and spun onto 384-well plates and processed for 53BP1 and □-H2AX staining (see Materials and Methods). The staining patterns of 53BP1 and □-H2AX in the presence and absence of ETP-induced DNA damage were similar to those seen in immune cell lines (Fig. 3a). Freshly isolated and frozen cell isolates yielded identical results.

**Fig. 3).**
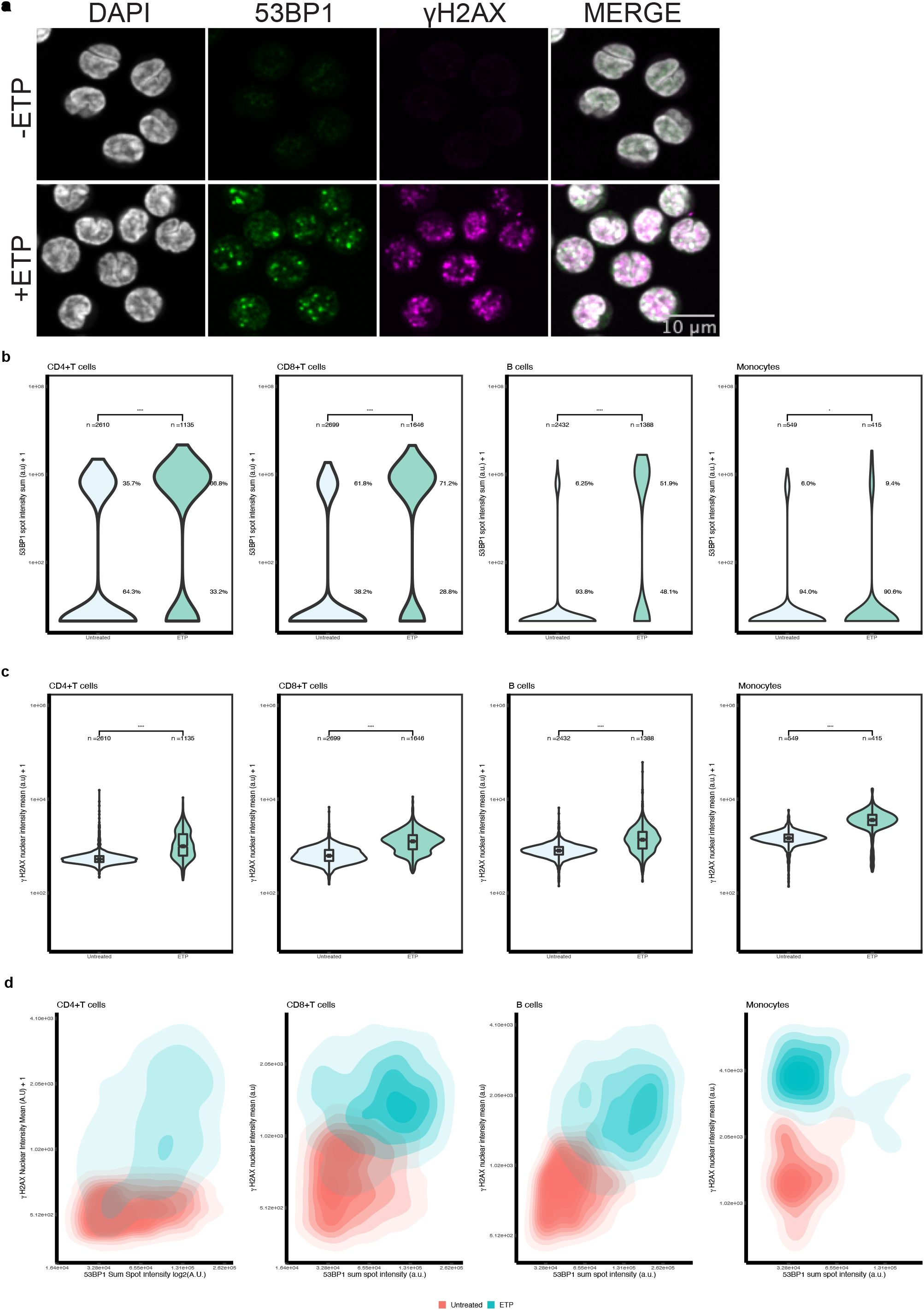
Detection of DNA damage in CD4^+^T, CD8^+^T, B cells and monocytes using HTI. **a)** CD4^+^T cells were either left untreated or treated with 30 µM ETP for 1.5 hrs before being subjected to HTI. DNA damage sites are indicated by formation of 53BP1 and *γ*-H2AX foci detected by specific antibodies. Scale bar: 10 µm. **b)** Distribution of 53BP1 sum spot intensity in DMSO- or ETP-treated cells. ****: p < 0.0001. **c)** Distribution of *γ*-H2AX mean nuclear intensity in DMSO- or ETP-treated immune cells. ****: p < 0.0001 using Mann-Whitney-Wilcoxon test. Number of analyzed cells (n) indicated for each sample. The values are representative of at least 3 experiments. Boxes represent the first and third quartiles of the distribution. The line represents the median. The whiskers represent the highest and lowest data points within 1.5 times the interquartile range above the upper quartile and below the lower quartile. **d)** Two-dimensional kernel density contour plots of the correlation between 53BP1 sum spot intensity and □-H2AX mean nuclear intensity for both ETP-treated (blue) and untreated (red) immune cell lines for the dataset shown in panels B and C. N = 3 wells per sample with at least 185 cells per well.

Quantitation of integrated 53BP1 spot intensity per cell for CD4^+^T cells showed an increase in response to ETP treatment (median: 8.71 ± 6.23 e5 A.U.) as compared to untreated cells (median: 5.24 ± 2.98 e4 A.U.) (Wilcoxon test p-value < 2.2 e-16) (Fig. 3b). In untreated cells, only 35.7% of cells were 53BP1 positive as compared to 66.8% of ETP-treated cells (*X*^2^ = 306.43, p < 2.2 e-16). In ETP-treated cells, □-H2AX mean nuclear intensity (median: 993 ± 652 A.U.) increased compared to untreated cells (median: 527 ± 112 A.U.) (Wilcoxon test p-value < 2.2 e-16) (Fig. 3c). Similarly, 47.3% of ETP-treated cells that had a nuclear intensity classified as positive-□-H2AX whereas only 33.3% showed H2AX positivity in untreated cells (Fig. 3c; *X*^2^ = 446.79, p-value <2.2 e-16).

We similarly detected DNA damage in all other primary immune cells. In CD8+T cells integrated 53BP1 spot intensity per cell increased from 4.58 ± 2.27 e4 A.U. to 8.36 ± 5.93 e4 A.U. in ETP-treated cells (Fig. 3b; Wilcoxon test, p < 2.2 e-16). We also found 69.3% of the cells in the ETP-treated condition were positive for 53BP1 as compared to 23.7% in the untreated condition (*X*^2^ = 875.77, p < 2.2 e-16). Furthermore, there was a significant increase in □-H2AX mean nuclear intensity in ETP-treated cells compared to untreated cells (median: 1266 ± 641 A.U. vs. 623 ± 244 A.U.; Wilcoxon test p-value p < 2.2 e-16) (Fig. 3c). Similarly, 13.5% of ETP-treated cells that had a nuclear intensity classified as positive-□-H2AX using a threshold of one SD above the mean □-H2AX nuclear intensity of the negative control, whereas only 0.2% of untreated cells were positive for H2AX (Fig. 3c; *X*^2^ = 357.51, p-value < 2.2 e-16).

B cells showed a significant increase in integrated 53BP1 spot intensity per cell upon ETP-treatment (median: 1.33 ± 0.85 e5 A.U.) when comparing to untreated cells (median: 4.76 ± 1.84 e4 A.U.; Wilcoxon test, p < 2.2 e-16) (Fig. 3b). Moreover, 51.9% of cells were 53BP1 foci positive in ETP treated cells compared to 6.25% in untreated cells (*X*^2^ = 1041.5, p < 2.2 e-16). We also observed a significant increase in □-H2AX nuclear intensity of ETP-treated B cells (median: 1368 ± 790 A.U.) when compared to untreated B cells (median: 801 ± 201 A.U., Wilcoxon test p < 2.2 e-16) (Fig. 3c). Similarly, 61.5% of ETP-treated cells that had a nuclear intensity classified as positive-□-H2AX compared to only 10.2% of untreated control cells (Fig. 3c, *X*^2^ = 584.03, p-value < 2.2 e-16).

For monocytes, we found a significant increase in integrated 53BP1 spot intensity per cell in response to ETP-induced DNA lesion (median: 5.00 ± 2.68 e4 A.U.) as compared to untreated control (median: 3.72 ± 0.99 e4 A.U.; Wilcoxon test p-value p = 3.7 e-3) (Fig. 3b). The number of cells positive for 53BP1 was not statistically different between control and ETP treated cells (6.0% vs 9.4%; *X*^2^ = 3.4476, *p* = 0.06). There was also a significant increase in *γ*-H2AX nuclear intensity when we compared ETP-treated cells (median: 3608 ± 1361 A.U.) to untreated cells (median: 1484 ± 382 A.U.; Wilcoxon test p < 2.2 e-16) (Fig. 3c). Similarly, 89.9% ETP-treated cells that had a nuclear intensity classified as positive-□-H2AX using a threshold of one SD above the mean □-H2AX nuclear intensity of the negative control, compared to only 10% of untreated cells (Fig. 3c, *X*^2^ = 607.38; p-value < 2.2 e-16).

Two-dimensional kernel density mapping confirmed the observed responses in all cell types (Fig. 3d) and the diagonal shift between control and ETP treated samples indicated that both DDR markers were simultaneously induced in individual cells. Taken together, these results demonstrate that a wide range of immune cells are amenable to quantitative interrogation of their DNA damage status in a high-throughput imaging format and that the extent of DNA damage can be sensitively measured in this assay.

### Inter-assay variability in CD4^+^T cells

In order to assess the robustness of HiIDDD, and to test whether this pipeline can be used to compare different biological samples in multiple batches, we determined the technical variability of our HTI pipeline between samples obtained from a single donor. To do so, we used three samples of frozen CD4^+^T cells from a single donor that were independently thawed, IF stained, imaged, and analyzed in different experimental batches (i.e., on different days). Cells were either left untreated, treated with DMSO as a vehicle, or treated with 30 µM ETP for 1.5 hrs. The 53BP1 integrated spot intensity per cell (Fig. 4a) and □-H2AX mean nuclear intensity per cell (Fig. 4b) were quantitated for each replicate and the variability of the biological replicates for each treatment group was analyzed via ANOVA. As expected ETP treatment significantly increased 53BP1 spot intensity (mean = 1.36 ± 1.04 e5) compared to DMSO vehicle control (mean = 2.17 ± 4.24 e4) and untreated cells (mean = 0.9171 ± 2.46 e4) (Fig. 4a, p = 1.9e-4). We similarly observed a modest increase in □-H2AX nuclear intensity when ETP-treated cells (mean = 2248 ± 2304) were compared to DMSO controls (mean = 811 ± 2174) or to untreated cells (mean = 770 ± 2678) (Fig. 4b, p = 0.011). No statistically significant differences were observed between individual replicates for 53BP1 spot intensity (p = 0.99) nor for □-H2AX nuclear intensity (p = 0.93). These results demonstrate that the HTI IF assay is robust and shows negligible batch to batch, or technical, variability.

**Fig. 4).**
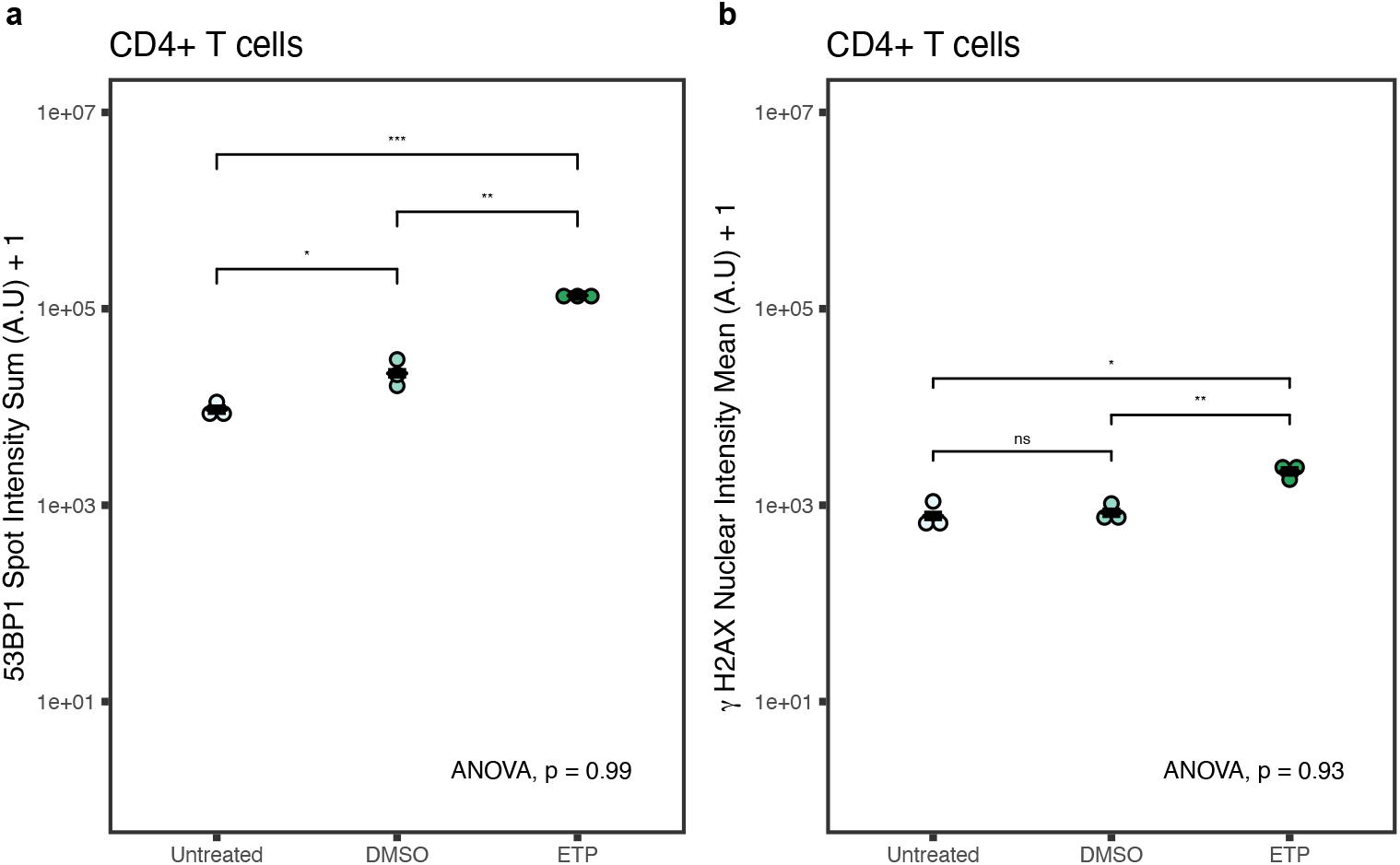
Inter-assay variability. **a)** Distribution of 53BP1 sum spot intensity in untreated, DMSO-treated, and ETP-treated CD4^+^ T cells from the same donor. Each dot represents the mean value of a treatment replicate. Black line represents mean integrated 53BP1 spot intensity and □-H2AX mean nuclear intensity *: p < 0.05, **: p < 0.01, ***: p < 0.001. ANOVA=0.99. **b)** Distribution of *γ*-H2AX nuclear intensity in untreated, DMSO-treated, and ETP-treated CD4^+^ T cells from the same donor. Each dot represents the mean *γ*-H2AX nuclear intensity for each treatment replicate. Black line represents the mean □-H2AX nuclear intensity for all three replicates. *: p < 0.05, ns: p>0.05. ANOVA=0.93. Statistics calculated using Mann-Whitney-Wilcoxon test and one-way ANOVA. N= 1776-14059 cells per sample.

### Batch processing of primary human immune cells

The intent of developing a high-throughput imaging assay for detection of DNA damage is its use in analyzing large sample numbers such as from archived materials that are part of longitudinal studies. This application would require batch processing of samples. To demonstrate scalability of HiIDDD, three primary immune cell types from 10 different donors were processed simultaneously (Supplementary Fig. 2). Cells, either untreated or treated with 30 µM ETP ex-vivo for 1.5 hrs, were processed in parallel and seeded in triplicate wells on a 384-well plate for imaging (see Materials and Methods). Elevated DNA damage, as indicated by an increase in 53BP1 and □-H2AX staining intensity, was detected in all samples to similar extents as in single processed samples (Fig. 5a, b), demonstrating scalability of the assay.

**Fig. 5).**
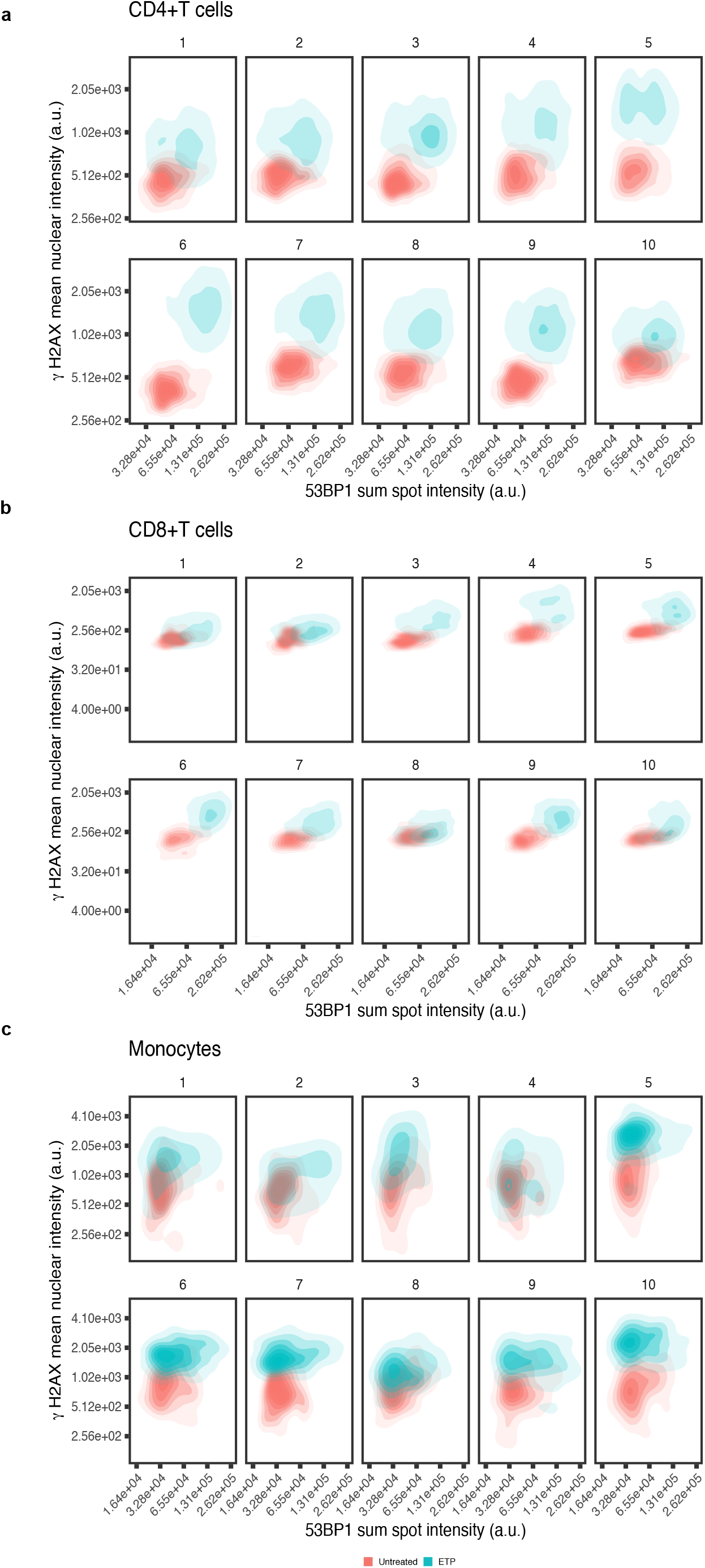
Batch processing of human primary immune cells for analysis of 53BP1 spot intensity and *γ*-H2AX nuclear intensity. CD4^+^T, CD8^+^T, and monocytes of 10 different donors were processed simultaneously, left untreated or treated with 30 µM ETP for 1.5 hrs and imaged in high-throughput format. 53BP1 sum spot intensity vs. *γ*-H2AX mean nuclear intensity were blotted for each donor for **a**) CD4^+^T cells, **b**) CD8^+^T cells **c**) monocytes. N= 663-5693 cells per sample.

Within each of the ten samples from different donors, the integrated 53BP1 spot intensity varied typically over a ∼5-fold range for all cell types. Similarly, □-H2AX mean nuclear intensity values in untreated cells were consistently distributed over an ∼8-10 fold range across samples from different donors, in line with values observed in samples from individuals (Fig. 5a-c). Among the samples from the ten donors, the integrated 53BP1 median spot intensity showed limited variation of ∼2-fold in CD4^+^T cells, ∼ 2-fold in CD8^+^T cells, and ∼ 1.5-fold in monocytes (Fig. 5A-C). Similarly, for □-H2AX median values varied between samples by ∼ 1.5-fold in CD4^+^T cells, ∼ 1.5-fold in CD8^+^T cells, and ∼ 2.5-fold in untreated monocytes (Fig. 5a-c).

We next challenged all ten samples by ETP-treatment to probe for potential differences in the extent to which the DDR is induced in individual donors. Increased 53BP1 and □-H2AX signals were detected in all samples and cell types (Fig. 5a-c). The distribution for both 53BP1 and □-H2AX broadened somewhat within each sample but largely stayed within a ∼6-fold range for 53BP1 and ∼10fold for □-H2AX (Fig. 5a-c). Comparing amongst the ten donor samples after ETP-treatment similar variation of ∼2-fold was observed for both 53BP1 median spot intensity and □-H2AX median values (Fig. 5a-c). The observed diagonal shift of 53BP1 and □-H2AX signal between untreated and treated cells in the dual parameter analysis also confirmed a concomitant increase in both damage markers in individual cells of all samples upon ETP treatment (Fig. 5a-c). No clear correlation between age or sex of the donor and baseline DNA damage was observed amongst the 10 samples derived from donors aged 30 to 85 (Fig. 5a-c). Interestingly, the extent of the response to ETP-treatment, however, appeared to differ between samples, with donors 1, 2, 5, and 10 showing less pronounced increases in DNA damage markers upon ETP treatment compared to the other donors (Fig. 5a-c). Importantly, the differences in response were seen concomitantly in all cell types of each individual donor. The coordinated response in multiple immune cells from the same donor points to inherent differences in the DNA damage response between individuals.

## Discussion

We have here developed HiIDDD, an imaging-based method for the detection of DNA damage in a wide variety of primary immune cells in a high-throughput format. HiIDDD is based on an optimized high-throughput imaging pipeline using a combination of both automated and manual techniques to maximize cell recovery following IF staining. We show that this assay is sensitive, robust and can be performed in a batch-processing format to assess both baseline DNA damage levels as well as DDR induction in primary immune cells.

HiIDDD relies on the detection of nuclear signals of the two well established DNA damage markers 53BP1 and □-H2AX ^7,8^. The method described here extends current approaches, such as cytometric analysis and traditional IF of H2AX phosphorylation status, that have previously been used to assess DNA damage^5,26–29^, in that it generates single cell data and does so in a high-throughput format suitable for large-scale studies. The sensitivity of 53BP1 and □-H2AX detection was comparable to standard low-throughput microscopy assays and induction of DNA damage by ETP resulted in a robust increase in signal for both 53BP1 and □-H2AX. The two measured signals are qualitatively distinct, and complementary, in that □-H2AX measures the de-novo formation of the epitope through phosphorylation of Ser139 on H2AX, whereas the 53BP1 signal measures the accumulation of the protein at sites of DNA damage. To capture these different properties of the two markers, mean nuclear intensity was measured for □-H2AX, but for 53BP1 the intensity of the signal in nuclear foci was detected. The bi-modal distribution of the 53BP1 signal reflects the presence of foci, albeit typically smaller in size, in undamaged cells. Reassuringly, the two DNA damage markers yielded corresponding responses with most cells exhibiting an increasing in both markers upon DNA damage. Importantly for an assay designed for high-throughput applications, we show low variability among cells from the same individual processed in different experimental batches, and high concordance in both basal and induced levels of the DDR in different cell types from the same individual. These observations indicate that HiIDDD is suitable for processing and analysis of large cohorts of samples in multiple separate experimental batches. Moreover, we find that our assay yields comparable results regardless of the use of fresh or cryopreserved primary immune cells, making it potentially suitable for interrogation of collections of archived materials.

While HiIDDD has been developed and optimize for the detection of DNA damage, the conditions used can similarly be applied to the detection of any cellular marker of choice. An important step in the detection pipeline is the efficient immobilization of immune cells on the imaging substrate, which has been a notoriously challenging aspect of imaging of immune cells. We describe here optimized methods to ensure sustained adherence of immune cells on the imaging substrate throughout the detection protocol by the combined application of a pre- and post-centrifugation fixation step to maximize the number of immobilized cells and the use of sponge evacuation for aspiration of fluids during all incubation steps rather than vacuum-mediated aspirations (see Methods). Using these steps, we typically find substantial retention of cells through the IF staining protocol, yielding enough cells for high-throughput imaging and collection of sufficiently large single-cell datasets for statistical analysis.

HiIDDD enables batch processing of relatively large sample sets. As proof of principle, we analyzed baseline DNA damage and DDR response capacity for 30 samples from ten donors using parallel processing. The total parallel processing of all samples time-divided into sample preparation, IF, and imaging, requires about 3 days with the most time-intensive steps being sample preparation and the manual fixation of seeded cells. In our experimental setup (see Materials and Methods) the imaging time per sample using three technical replicates is ∼ 3 min. Processing time could be reduced by use of more extensive robotics for liquid handling during fixation and DNA damage induction or the use of X-ray or gamma-irradiation of entire 384-well plates without the need for drug treatment and washing. The versatility and capacity for high-throughput make the HiIDDD imaging pipeline useful for basic research applications that involve larger sample sets, for longitudinal or natural history studies, and also for clinical applications such as the assessment of immunotherapy responses.

One limitation in our current work is the inherent variability of 53BP1 and □-H2AX foci morphology, and of the signal intensity amongst different immune cell types (Fig. 2, 3, 5). A potential strategy to gain more precision is to increase the utility of our detection method is the use of additional biomarkers. The two current markers could be combined with other DNA damage markers such as components of the MRE11 complex^37^ or downstream effectors such as CHEK1 or CHEK2^10,37–39^. Alternatively, non-DNA damage related indicators such as epigenetic markers, could be included in a multiplexed assay. By using multi-dimensional analysis, it should be possible to more accurately cluster single-cell data to reflect DNA damage status.

In summary, we have developed a high-throughput imaging pipeline to detect and quantitatively measure DNA damage in primary human immune cells. Our assay can detect both baseline levels of DNA damage between samples in addition to DNA damage response capacity. The assay is scalable and sensitive to differences in DNA damage response between individuals and represents a robust and easy strategy to measure DNA damage response in immune cells in large sample sets.

## Materials and Methods

### Cell culture

The Jurkat (Clone E6-1, TIB-152) and THP-1 (TIB-202) cell lines were purchased from ATCC. BJAB cells were a kind gift of Ranjan Sen Lab (National Institute of Aging/NIH). All cell lines were cultured in high glucose Dulbecco’s modified eagle’s medium (DMEM, ATCC, cat. number 30-2002), 10% fetal bovine serum (Gibco, cat. number A3410), and 1% penicillin/streptomycin (Gibco, cat. number 15140122). All cell lines were grown at 37°C and 5% CO_2_. Cells were maintained according to vendor’s recommendations and kept at low passage numbers (≤ 6 passages).

### Primary cells processing

Study samples were derived from Genetic and Epigenetic Signatures of Translational Aging Laboratory Testing (GESTALT) participants (n=5) and normal donors (n=5) ages 20-85 years. Participants enrolled in GESTALT (NIA protocol# 15-AG-0063) are volunteers who at the time of study enrollment were “healthy” based on eligibility criteria^40^. Normal donors are volunteers who donated apheresis material through the Cytapheresis of Volunteer Donors protocol (NIA protocol# 03-AG-N316). All individuals were consented for their donations and protocols have been reviewed by the NIH institutional review board.

PBMCs were isolated from apheresis packs using Ficoll-Paque Plus (GE Healthcare, Piscataway, NJ, USA cat. number 17-1440-03) density gradient centrifugation. CD4^+^ T cells, CD8^+^ T cells, B cells, and monocytes were separated from freshly isolated PBMCs using immunomagnetic negative selection cell isolation kits (EasySep Human Cell Enrichment Kits; CD4^+^ T cells: cat. number 19052; CD8^+^ T cells: cat. number 19053; B cells: cat. number 19054 and monocytes, without CD16 depletion: cat# 19058; StemCell Technologies, Vancouver, BC, Canada) using a fully automated cell separator (RoboSep, StemCell Technologies) according to the manufacturer’s protocol.

CD4^+^ T cells, CD8^+^ T cells, B cells, and monocytes were cryopreserved in freezing medium (FBS from Life Technologies, cat. number 10437028 and 15% DMSO from Sigma Aldrich, cat. number D2650) in a freezing container overnight at −80° C and transferred to below −140° C liquid nitrogen freezer until use. Cryopreserved cells were thawed using RPMI 1640 containing 10% fetal calf serum.

### Etoposide treatment

Cell lines and isolated primary human immune cells were treated with etoposide (ETP, Sigma-Aldrich, cat. number E1383) at the indicated concentrations from a 50 mM stock that was prepared from a lyophilized powder reconstituted in DMSO (Sigma-Aldrich, cat. number 20-139). For treatment, cells were seeded at 1.25 × 10^6^ cells/ml in 6-well flat bottom cell culture plates (Corning, cat. number 07-200-83). The incubation time with etoposide or DMSO for all experimental conditions was 1.5 hrs at 37°C and 5% CO_2_.

### Semi-automated sample preparation and immunofluorescence (IF) staining in 384-well plates

For the DNA damage detection assay, cells were washed once with D-PBS (R&D Systems, cat. number 30250) and fixed with 4% paraformaldehyde (PFA, Electron Microscopy Sciences, cat. number 50-259-96) in D-PBS for 20 min at RT. Cells were washed twice with D-PBS and then resuspended in D-PBS before being seeded on 384-well plates with poly-D-lysine coating (PerkinElmer CellCarrier Ultra 384, cat. number 6057500). Plates were pre-warmed at 37°C for 20 min prior to cell seeding. Cells were routinely stored for up to 72 hrs at 4°C prior to seeding on plates. Following a centrifugation step (400 g, 5 min, at RT), cells were crosslinked to plates using 4% PFA for 20 min before washing with D-PBS. All washes of 384-well plates were completed using an automated BlueWasher liquid handler (BlueCatBio) with settings optimized for low-adhering cells. To wash cells, 40-60µl/well of D-PBS was dispensed at a pressure level of 1 with auto-prime and staccato dispense cadence enabled. Dispense position offset was adjusted to –0.5. Liquid was gently aspirated by sponge evacuation by placing 384-well plates over a standard utility sponge overlayed with a stainless-steel mesh screen. Pressure was lightly applied to the back of the plate to promote evacuation of wash liquid. This process was repeated 2-3 times. Cells were permeabilized with 0.5% Triton X-100 (Thermo Scientific, cat. number 85111) diluted in D-PBS for no more than 8 min before washing with D-PBS. Primary and secondary antibodies were suspended in antibody buffer containing 1% bovine serum albumin (BSA, Sigma Aldrich, cat. Number A9418) and 0.1% Triton X-100 in D-PBS. Primary antibodies were incubated overnight (8-14 hrs) at 4°C, and secondary antibodies were incubated for 1 hr at RT. Cells were washed twice in D-PBS after each antibody incubation period.

The following antibodies were used: Primary rabbit anti-53BP1 (1:1200, Novus Biological, cat. number NB100-304) and primary mouse monoclonal anti-□-H2AX (1:1200, Sigma-Aldrich, cat. number 05-636). 4’,6-diamidino-2-phenylindole (DAPI) (10µg/ml, Biotium, cat. number 40043) was used for nuclear staining. Labelled secondary antibodies were goat anti-mouse AlexaFluor488 (Invitrogen, cat. number A28175) or donkey anti-rabbit AlexaFluor647 (Invitrogen, cat. number A-31573). Both secondary antibodies were used at a concentration of 1:1000.

### High throughput image acquisition and analysis

Cells were imaged using a CV7000 (Yokogawa) high throughput spinning disk confocal microscope. Images were acquired using a 60X water-immersion lens (NA 1.2) and 2 sCMOS cameras (2560 × 2160 px, physical pixel size 6.5 microns) using 2× pixel binning (pixel size 216 nm). For excitation of the fluorophores, three laser lines (DAPI: 405 nm, Alexa488: 488 nm, and Alexa647: 640 nm), a fixed excitation dichroic mirror (405/488/561/640 nm), a fixed emission mirror (568 nm), and matched emission bandpass filters (DAPI: 445/45nm, Alexa488: 525/50 nm, and Alexa647: 676/29 nm) were used in separate exposures for each fluorophore. All wells were imaged using 9 randomly selected fields of view (FOVs). Each experiment included at least 3 technical replicates (wells on the same plate) per condition. Typically, approximately 1-2 × 10^3^ cells were analyzed per well. Acquired images were processed using Columbus high-content image analysis software (PerkinElmer, version 2.8.1 and 2.9.1). The analysis pipeline first identifies and segments nuclei using the DAPI image. Within identified and segmented nuclei, 53BP1 and □-H2AX foci were then detected using the Columbus spot-detection algorithm with low stringency detection parameters. A supervised machine learning algorithm (Fisher Linear Classifier) was used to classify 53BP1 and □-H2AX foci from background signals. The pre-trained model was fine-tuned by retraining to a dataset from each cell type to achieve robust and accurate classification of 53BP1 and □-H2AX □-H2AX foci. To reflect the distinct staining patterns of the two markers integrated spot intensity was measured for 53BP1 and mean nuclear intensity for □-H2AX. The Columbus analysis sequence (.aas) files are available upon request.

Acquired images from the CV7000 were saved as .tif files and processed in Fiji, where only changes to adjust brightness and contrast on entire images were applied uniformly to all images of the same channel in a single figure panel.

### Statistical analysis

Single-cell data was exported as tabular text files and further analyzed and plotted using R (R Core Team, 2019) and RStudio 1.3 (RStudio Team, 2020) using the tidyverse package ^41^. All p-values for violin-boxplots of etoposide treated, untreated, and/or vehicle control samples were calculated using the Mann-Whitney-Wilcoxon test for pairwise comparisons, and the ANOVA test for multiple comparisons. Dot plots showing intra-assay variability between biological replicates in each condition also used Mann-Whitney-Wilcoxon and ANOVA to generate p-values. Unless indicated otherwise, experiments were performed at least as biological triplicates. The R analysis scripts used to generate the plots in the figures are deposited at https://github.com/gallantk72/hti-ddr.

## Acknowledgments

Research in the Misteli lab and at the NCI HiTIF is supported by funding from the Intramural Research Program of the National Institutes of Health (NIH), National Cancer Institute, and Center for Cancer Research (ZIAABCO10389). KG is an NCI iCURE Scholar.

## Author Contributions

KG: Conceptualization, Methodology, Software, Validation, Formal Analysis, Data curation, Data visualization, Investigation, Writing-original draft preparation, Writing – review and editing

AB: Methodology, Resources, Writing- original draft preparation, Writing-review and editing

MK: Resources, Writing- original draft preparation, Writing- review and editing

AL: Resources, Writing – review and editing

LF: Conceptualization, Resources, Writing- original draft preparation, Writing- review and editing, Supervision

GP: Conceptualization, Methodology, Software, Formal Analysis, Writing-original draft preparation, Writing- review and editing, Supervision

TM: Conceptualization, Methodology, Funding, Formal Analysis, Investigation Writing-original draft preparation, Writing – review and editing, Supervision

## Supplementary Figures

**Supplementary Fig. 1) Detection of 53BP1 spot intensity in immune cells**. 53BP1 positive cells were characterized in each treatment group for **a)** immune cell lines (Jurkat, BJAB, and THP-1) and **b)** CD4^+^T cells, CD8+T cells, monocytes, and B cells. The same samples as in figures 2 and 3 are used.

**Supplementary Fig. 2) List of donors used in batch processing assay. a)** For each listed donor in Figure 5, donors were identified by age and sex.

