## Supplementary Information for "HiIDDD: A high-throughput imaging pipeline for the quantitative detection of DNA damage in primary human immune cells"

**a**

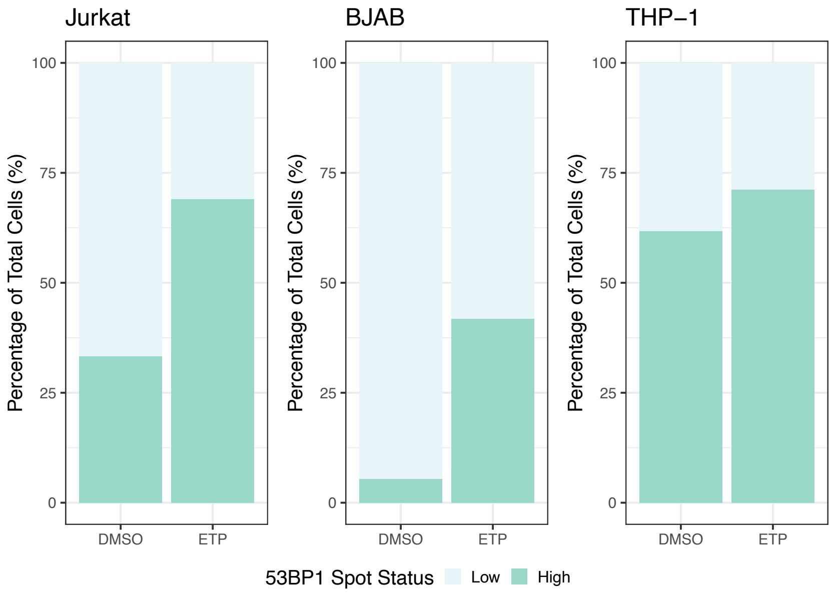
Jurkat
BJAB
THP−1
Percentage of Total Cells (%)
100
75
50
25
0
DMSO
ETP
53BP1 Spot Status
Low
High

**b**

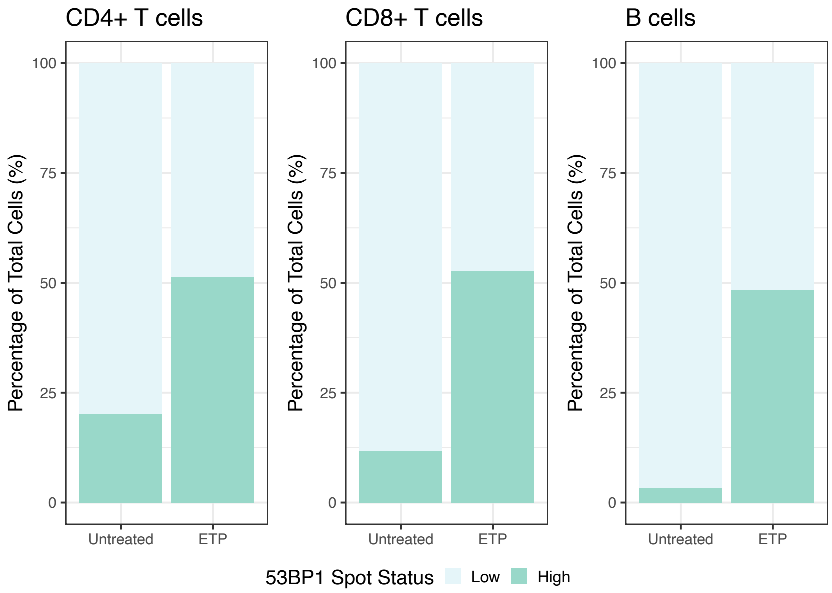
CD4+ T cells
CD8+ T cells
B cells
Percentage of Total Cells (%)
100
75
50
25
0
Untreated
ETP
53BP1 Spot Status
Low
High

**a**

| # | Donor | Age | Sex |
|---|---|---|---|
| 1 | ND791 | 30 | F |
| 2 | ND782 | 31 | F |
| 3 | Gt016 | 42 | F |
| 4 | Gt024 | 47 | M |
| 5 | Gt030 | 49 | M |
| 6 | ND550 | 54 | F |
| 7 | ND797 | 58 | F |
| 8 | Gt004 | 62 | M |
| 9 | ND334 | 67 | F |
| 10 | Gt029 | 85 | M |

**Supplementary Figures**

**Supplementary Fig. 1) Detection of 53BP1 spot intensity in immune cells.** 53BP1 positive cells were characterized in each treatment group for **A)** immune cell lines (Jurkat, BJAB, and THP-1) and **B)** $CD4^{+}$T cells, CD8+T cells, monocytes, and B cells. The same samples as in figures 2 and 3 are used.

**Supplementary Fig. 2) List of donors used in batch processing assay. A)** For each listed donor in Figure 5, donors were identified by age and sex.
